# Cenozoic evolutionary history obscures the Mesozoic origins of acanthopterygian fishes

**DOI:** 10.1101/2024.09.30.615987

**Authors:** Chase D. Brownstein, Alex Dornburg, Thomas J. Near

## Abstract

Sister lineage comparisons provide a valuable tool for understanding evolutionary origins of species-rich clades. *Percomorpha*, comprising over 18,900 species, represents one of the most species-rich vertebrate clades. However, the phylogenetic resolution of its sister lineage remains unclear, obscuring whether contrasts in histories of diversification provide insights into the factors that gave rise to this clade’s diversity. Using 887 ultraconserved element loci and Sanger-sequenced nuclear genes, we resolve the phylogenetic relationships of the three closest relatives of *Percomorpha-*the roughies, flashlightfishes, porcupinefishes and fangtooths (*Trachichthyiformes*), the squirrelfishes and soldierfishes (*Holocentridae*), and the whalefishes, bigscales, and alfonsinos (*Berycoidei*)-and the placement of percomorphs among them. Contrary to expectations from the fossil record, we demonstrate that living lineages of *Berycoidei*, *Holocentridae*, and *Trachichthyiformes* all diversified after the Cretaceous-Paleogene mass extinction. Our findings show that multiple clades in *Trachichthyiformes* and *Berycoidei* independently colonized deep ocean habitats during the climatically unstable Eocene and Oligocene and shallow-water reefs during the extensive hotspot migration and faunal turnover of the Early Miocene. Due to their complex evolutionary history, the closest relatives of *Percomorpha* are not ideal for understanding the origins of this exceptionally species-rich clade.

## Introduction

The last 100 million years of Earth History have witnessed dramatic changes in the composition of marine biotas. In particular, the extant diversity of marine fishes has been fundamentally impacted by the rise to dominance of the spiny-rayed fish clade *Acanthomorpha* (Friedman 2010, 2022; Near et al. 2012, 2013; Betancur-R et al. 2013, 2017; Alfaro et al. 2018; Hughes et al. 2018; Ghezelayagh et al. 2022), which contains nearly 20,000 species including iconic lineages such as seahorses, cods, gobies, wrasses, pufferfishes, anglerfishes, perches, and flatfishes (Eschmeyer’s Catalog of Fishes | California Academy of Sciences n.d.). Acanthomorph evolution provides an excellent system for studying rapid lineage diversification, but inferences of the ages and ancestral morphologies of its most species-rich subclades is hampered by disagreement around the phylogenetic relationships of some of its oldest branches (Miya et al. 2003; Near et al. 2012, 2013; Betancur-R et al. 2013, 2017; Dornburg et al. 2017; Hughes et al. 2018; Ghezelayagh et al. 2022). In particular, which of three relatively species-poor clades, *Berycoidei* (alfonsinos, bigscales, and whalefishes), *Holocentridae* (squirrelfishes and soldierfishes), and *Trachichthyiformes* (roughies, fangtooths, flashlightfishes, spinyfins, and porcupinefishes), is the living sister to the lion’s share of acanthomorph diversity in *Percomorpha*, remains unclear. Although these lineages are consistently resolved together with percomorphs in a clade called *Acanthopterygii*, nearly every possible arrangement of *Berycoidei, Holocentridae, Percomorpha*, *Trachichthyiformes* has been resolved across phylogenies inferred using morphology (Stiassny and Moore 1992), Sanger-sequenced mitochondrial and nuclear DNA sequences (Miya et al. 2003; Near et al. 2012, 2013; Betancur-R et al. 2013, 2017), and genome-scale DNA sequence datasets (Alfaro et al. 2018; Hughes et al. 2018; Musilova et al. 2019; Ghezelayagh et al. 2022).

This issue compounds the dearth of information available on the early evolution of acanthomorphs from the fossil record, which is notoriously poor for the interval when they initially diversified in the Cretaceous-Paleogene (Patterson 1993; Davesne et al. 2016; Friedman 2022; Friedman et al. 2023).

Several early acanthomorph fossils have variously been placed in all three of the potential sister lineages of *Percomorpha*, hampering their utility for both informing our understanding of the origins of acanthomorph body plans and calibrating the timescale of spiny-rayed fish evolution (Andrews et al. 2023). Whether the living representatives of the *Berycoidei*, *Holocentridae,* and *Trachichthyiformes* are also representative of ancient acanthomorph ecological and morphological diversification is also unclear, as different analyses have produced considerably different estimates of the ages of these lineages (Near et al. 2012, 2013; Betancur-R et al. 2013, 2017; Dornburg et al. 2015; Alfaro et al. 2018; Hughes et al. 2018; Ghezelayagh et al. 2022) and disputed fossil evidence suggests that these clades diversified in marine environments as early as the Late Cretaceous (Muséum national d’histoire naturelle (France) and naturelle (France) 1951; Patterson and Patterson 1967; Bardack and Bardack 1976; Patterson 1993; Patterson and White 1997; Friedman et al. 2023). Time-calibrated phylogenies produced for *Berycoidei*, *Holocentridae,* and *Trachichthyiformes* using both legacy markers (Near et al. 2012, 2013; Betancur-R et al. 2013) and genome-scale DNA sequence data (Hughes et al. 2018; Ghezelayagh et al. 2022) suggest post-Cretaceous origins for these crown clades. However, a species-rich timetree for all three of these lineages has yet to be generated.

Here, we reconstruct a phylogeny sampling all taxonomic families of non-percomorph *Acanthopterygii* to infer their evolutionary relationships and estimate their timescale of diversification. Using a combination of genome-scale sequence data and Sanger-sequenced nuclear genes, we resolve the living sister to percomorphs and show that all three of the potential percomorph sister clades have a history of recent diversifications into reef and deep-sea ecosystems. Our results exclude the possibility that Cretaceous fossils are members of crown *Berycoidei*, *Holocentridae,* and *Trachichthyiformes* and constrain the timescale of percomorph evolution and the emergence of acanthomorphs as the dominant clade of marine vertebrates.

## Methods

### (a) DNA Sequencing and Dataset Assembly

We combined ultraconserved element (UCE) sequences with Sanger-sequenced nuclear genes to produce a dataset spanning all taxonomic families and approximately 35% of named species in the clades *Berycoidei*, *Holocentridae*, and *Trachichthyiformes* sensu Near and Thacker (Near and Thacker 2024). Ultraconserved elements were compiled from published genomes available on Genbank and contigs from two published phylogenetic analyses of *Acanthomorpha* (Alfaro et al. 2018; Ghezelayagh et al. 2022). We extracted, processed, and combined UCEs from published genomes (Supplementary Table 1) with UCE contigs using phyluce v. 1.7.3 (Faircloth 2016) to produce a 75% complete dataset and tested and removed chimeric sequence data using CIAlign (Tumescheit et al. 2022). The final UCE dataset consisted of 879 UCEs collected for 44 individuals, including 35 species of *Berycoidei*, *Holocentridae*, and *Trachichthyiformes,* for all major family-level and subfamily-level clades (*Anomalopidae, Anoplogaster, Barbourisia, Cetomimidae, Diretmidae, Gibberichthys, Holocentrinae, Melamphaidae, Monocentridae, Myripristinae, Rondeletia, Stephanoberycidae, Trachichthyidae).*We Sanger-sequenced genes for this study using markers employed in previous studies of ray-finned fishes (Dornburg et al. 2012; Near et al. 2012, 2013) for the following genes: *enc1, glyt*, *myh*, *plagl2*, *ptr*, *rag1*, *sreb*, *zic*. New sequences were aligned with previously published sequences downloaded from the NCBI sequence repository Genbank. We ran preliminary maximum likelihood analyses on each of the Sanger-sequenced nuclear gene alignments to identify poor-quality and misidentified samples.

### (b) Maximum Likelihood Phylogenetic Analyses

A maximum likelihood phylogeny was inferred from the UCE dataset using IQ-TREE2 (Nguyen et al. 2015; Minh et al. 2020b). For all analyses, we used ModelFinder (Kalyaanamoorthy et al. 2017) to find best-fit models of molecular evolution for individual UCEs or partitions, and partition schemes were selected using PartitionFinder 2 (Lanfear et al. 2012, 2017) as implemented in IQ-TREE2. To assess the impact of partitioning and gene-tree species tree heterogeneity, we additionally conducted analyses where the UCEs were concatenated and treated as a single partition, and also generated gene trees from each locus for multispecies coalescent tree building in ASTRAL-III (Zhang et al. 2018). In all cases, 1000 ultrafast bootstrap replicates were used to assess nodal support. To refine our assessment of support values across nodes, we further calculated gene and site concordance factors using the concatenated single-partition topology and the gene trees (Minh et al. 2020a). Gene and site concordance factors provide information on the proportions of gene trees and sites that support a given node, and thus provide additional information on the drivers of varying levels of support across a tree topology (Minh et al. 2020a; Kück et al. 2022). We ran all analyses involving UCEs on the Yale High-Performance Computing Cluster McCleary.

We concatenated alignments of the Sanger-sequenced nuclear genes and ran an analysis in IQ-TREE (Trifinopoulos et al. 2016) using 1000 ultrafast bootstrap supports, Shimodaira– Hasegawa approximate likelihood ratio tests (Shimodaira and Hasegawa 1999) and approximate Bayes tests (Anisimova et al. 2011) to assess nodal support, and allowing for free rate heterogeneity. We inferred phylogenies using a concatenated dataset of the UCE and Sanger-sequence nuclear gene sequences with a single-partition IQ-TREE2 analysis on the Yale Computing Cluster with an ultrafast bootstrap generated over 1000 replicates to assess support. We also generated gene trees for the UCEs and each Sanger-sequenced nuclear gene to produce an ASTRAL-III species tree combining these data types.

### (c) Bayesian Phylogenetic Analysis and Time-Calibration

We estimated the timescale of divergences among *Percomorpha* and its closest relatives among acanthomorph fishes via a set of node-dated relaxed molecular analyses conducted in BEAST v 2.6.7. (Bouckaert et al. 2014, 2019) using a newly-compiled set of fossil calibrations (Supplementary Information). The affinities of many extinct species to *Berycoidei*, *Holocentridae*, *Trachichthyiformes*, and *Percomorpha* are controversial, and recent studies have found that key Late Cretaceous fossils show unclear affinities to these crown clades. In particular, putative global Cretaceous otolith records for *Beryx* and related forms (Schwarzhans et al. 2017) and holomorphic fossils of berycoids, holocentrids, and trachichthyiforms from Lagerstätten in Lebanon (Patterson and Patterson 1967) should be viewed with in the context of recent phylogenetic analyses of morphological data (Andrews et al. 2023). Because of this, we restricted our fossil calibration list to undisputed representatives of clades within *Berycoidei*, *Holocentridae*, *Trachichthyiformes*, and *Percomorpha*, and supplemented these calibrations with those for nodes in the closely related *Paracanthopterygii* that we use as outgroups in our phylogenetic analyses. Fossils used here as node calibrations include the fossil pan-holocentrid †*Iridopristis parrisi* from the earliest Paleocene of New Jersey (Andrews et al. 2023), the fossil velvet whalefish †*Miobarbourisia aomori* from the Middle to Late Miocene of Japan, and †*Scopelogadus* sp. from the Middle to Late Miocene of Russia (Nazarkin and Kotlyar 2020). We note that the phylogenies inferred from the UCE loci and the concatenated Sanger-sequenced nuclear genes are congruent in terms of in resolving both *Poromitra* and *Melamphaes* as reciprocally monophyletic and *Rondeletia* species as monophyletic relative to *Hispidoberyx ambagiosus*; however, analyses of combined datasets resolve each of these clades as non-monophyletic with very weak support. We believe this results from the lack of overlapping Sanger-sequenced nuclear gene data for species included in the UCE dataset. In our relaxed molecular clock analyses, we set priors that enforced the reciprocal monophyly of these lineages and allowed us to employ a fossil melamphaeid for a node-based calibration. Similarly, we constrained the position of *Anoplogaster* as sister to *Trachichthyidae*, as this relationship is obtained and supported by bootstrap and coalescent values of 100% and 1.0, respectively, in analyses using UCEs (Figures S1-S3, S6-S7) but weakly rejected when only Sanger-sequenced nuclear genes are used (Figure S5).

For the time-calibration analysis, we randomly subsampled all loci to produce three sets of 50 UCE loci and combined them with the sanger-sequenced data. We input a general time- reversable model of nucleotide evolution, a log-normal relaxed clock for the clock model, and the Fossilized Birth-Death (FBD) model as implemented in BEAST2 (Gavryushkina et al. 2016). We set the rho prior, which represents the proportion of living species sampled, to 0.36, following species counts from Eschmeyer’s Catalogue of Fishes (Eschmeyer’s Catalog of Fishes | California Academy of Sciences n.d.), and the diversification rate prior to 0.05 following a recent estimate of the background diversification rate of *Acanthomorpha* (Ghezelayagh et al. 2022). For the origin prior, we inputted an age of 145.0 Ma, the Jurassic-Cretaceous Boundary, with bounds of 161.5 Ma (the end of the Oxfordian Stage of the Jurassic), the age of the oldest defnite crown teleosts (Friedman 2022), and 93.9 Ma, which is when the youngest definite crown acanthomorphs appear in the fossil record (Newbrey et al. 2013; Davesne et al. 2014, 2016; Delbarre et al. 2016; Brownstein and Near 2024). For all node calibrations, we used lognormal priors and modified prior distributions such that 97.5% of the prior age in each case fell before the fossil age. We ran each BEAST analysis of the 50 UCE loci three times for 4.0 x 10^8^ generations with a 5.0 x 10^7^ pre-burnin, checked for convergence of the posteriors after 50% burnin in Tracer v 1.7 (Rambaut et al. 2018), and combined the last 25% of posterior trees from each run with subsampling every 5000 generations using LogCombiner v. 2.6.7 and TreeAnnotator v. 2.6.6 (Bouckaert et al. 2014, 2019) to produce a maximum clade credibility tree with median node heights.

### (d) Ancestral State Reconstructions and Trait-Associated Diversification Rates

We conducted an ancestral state reconstruction of habitat preference using the time-calibrated Bayesian phylogeny produced in BEAST 2.6.7 and habitat data from FishBase.se. Habitat preference was categorized as three states following previous studies (Miller et al. 2022; Horowitz et al. 2023): epipelagic (0-200 m), epipelagic-mesopelagic (0-1000 meters), and epipelagic-bathypelagic (0-1000 m or deeper). We used the R package phytools (Revell 2012) to conducted stochastic mapping over 1000 simulations and summarized the ancestral state reconstructions along a single tree. Next, we used Binary and multi-state dependent diversification rate analyses (BiSSE, MuSSE) as implemented in the R package diversitree (FitzJohn 2012) to test for associations between habitat, life history traits, and lineage diversification along the time-calibrated phylogeny. Using BiSSE and MuSSE, we tested three alternative coding schemes for the habitat trait, two traits relevant to the ontogeny and sexual dimorphism observed in some berycoids, and a reef association trait. Data on reef association were taken from Fishbase.se, and data on the other two traits were taken from the literature and the results of phylogenetic analyses in the current study and three previous ones (Johnson et al. 2009; Ghezelayagh et al. 2022; Kobyliansky et al. 2023). For the BiSSE analyses, we tested models with equal or variable speciation rates, and for the MuSSE analysis, we tested models differentially constraining transition, speciation, and extinction rates. Next, we conducted Markov chain Monte Carlo runs for 1.0 x 10^4^ generations on each best fit model and sampled every 100 generations. After burning 10% of the generations, we confirmed convergence of posterior values and plotted histograms of estimated diversification rates by state, along with mean values.

### (e) Nomenclatural Note

In this paper, our approach to the systematics and taxonomy of ray-finned fishes follow the conventions of the *PhyloCode* (Queiroz and Cantino 2020; Near and Thacker 2024). As such, we italicize all formal clade names (Thines et al. 2020).

## Results

### (a) Phylogenetic Relationships of Percomorpha and Its Closest Relatives

Our maximum likelihood phylogenies and Bayesian maximum clade credibility tree (Figure 1) of the UCE and the combined UCE and Sanger-sequenced nuclear gene datasets all resolve *Beryciformes* as containing *Holocentridae* and *Berycoidei* (Dornburg and Near 2021; Near and Thacker 2024) and a clade containing *Percomorpha* and *Beryciformes*. This result matches the results of previous analyses using UCEs (Alfaro et al. 2018; Ghezelayagh et al. 2022) anchored hybrid enrichment (AHE) loci (Dornburg et al. 2017), exons ((Hughes et al. 2018): figure S2), and whole genomes (Musilova et al. 2019), but is incongruent with phylogenies inferred using Sanger-sequenced mitochondrial and nuclear genes, which alternatively resolve *Holocentridae* (Betancur-R et al. 2013, 2017; Hughes et al. 2018) or a clade containing *Berycoidei, Holocentridae,* and *Trachichthyiformes* (Near et al. 2012, 2013) as the sister lineage of *Percomorpha.* Notably, the maximum likelihood phylogeny inferred from the Sanger-sequenced nuclear genes weakly resolves *Holocentridae* and *Percomorpha* as sister lineages (Figure S5; bootstrap value = 83%). An examination of gene and site concordance factors calculated using the ASTRAL-III and concatenated single-partition trees generated from the UCE-only dataset reveals the monophyly of both *Beryciformes* and the clade containing *Beryciformes* and *Percomorpha* are supported by relatively small proportions of sites and UCE loci. Although coalescent and ultrafast bootstrap supports for these clades are consistently 1.0 and 100% among the inferences using the UCE-only dataset (Figure 1; Figure 2; Figures S1-S3), these results highlight the earliest divergences among acanthopterygians as a region of phylogenetic discordance that is resolved only using genome-scale data (Dornburg et al. 2017; Ghezelayagh et al. 2022). Surprisingly, this discordance does not seem to result from rapid successive divergences, as multi-million-year periods appear to separate these lineages (Figure 1). Thus, molecular data favors a sister-lineage relationship of percomorphs and beryciforms with *Trachichthyiformes* resolved as the sister lineage to this clade within *Acanthopterygii* (Near and Thacker 2024). Further, this level of discordance does not affect ingroup relationships in *Berycoidei*, *Holocentridae*, and *Trachichthyiformes*, which are consistently resolved across the different phylogenies inferred using UCEs and Sanger-sequenced nuclear genes (Figures S1-S3, S5-S7).

**Figure 1.**
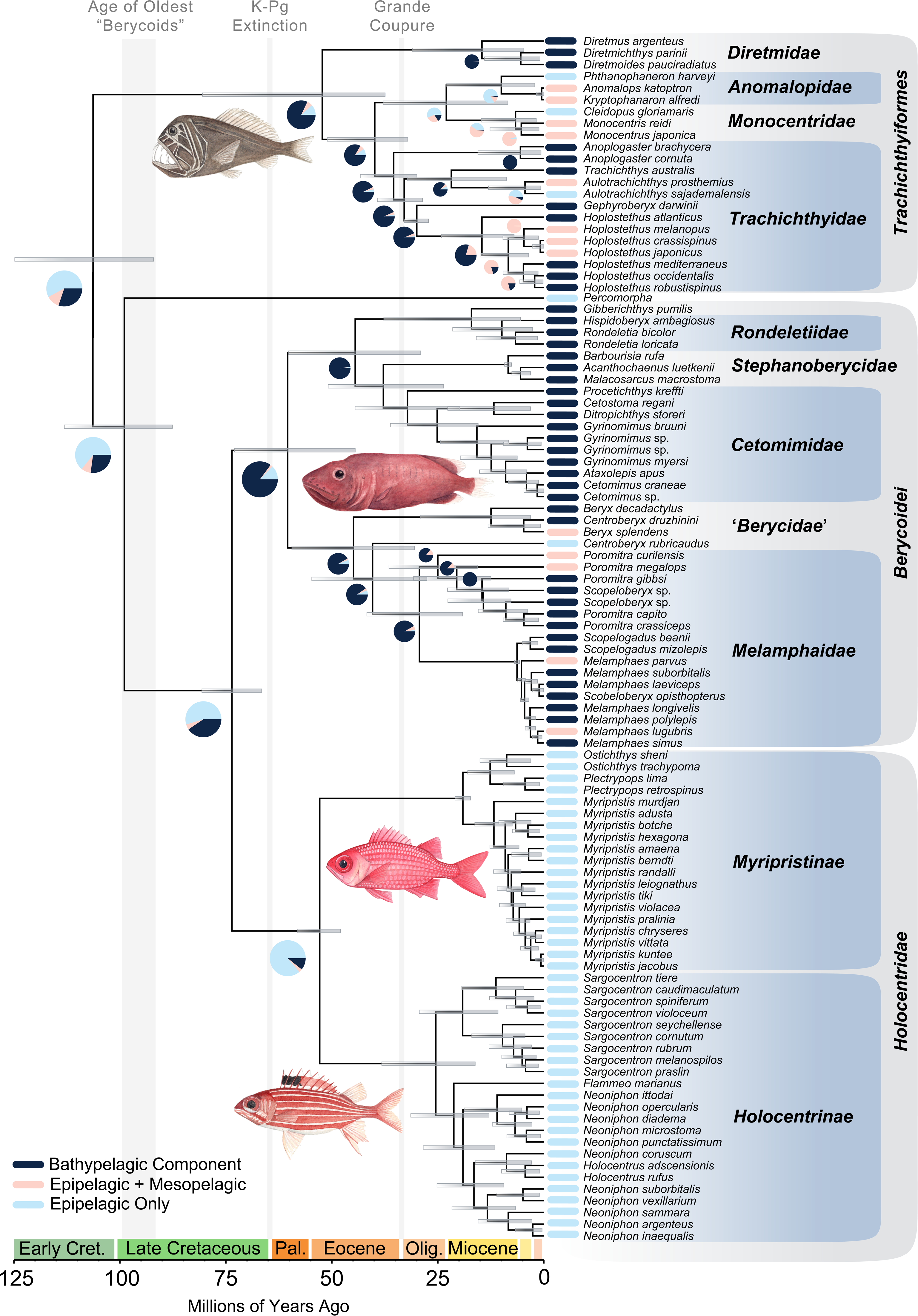
The Timescale of Acanthopterygian Evolution. The figure shows the node-dated Bayesian maximum clade credibility tree with median node heights resulting from multiple independent runs of three sets of 50 UCE loci and Sanger-sequenced nuclear genes. Dashed lines indicate clades in *Melamphaidae* that were enforced as monophyletic (see Methods). Bars at nodes indicate 95% highest posterior density intervals, and pie charts indicate inferred ancestral habitat states at nodes where the favored state is reconstructed with <80% support. Successive nodes have the same state as deeper ones unless denoted. Colors at tips indicate tip states. Abbreviations: Pal., Paleocene; Oli., Oligocene. Illustrations are by Julie Johnson.

**Figure 2.**
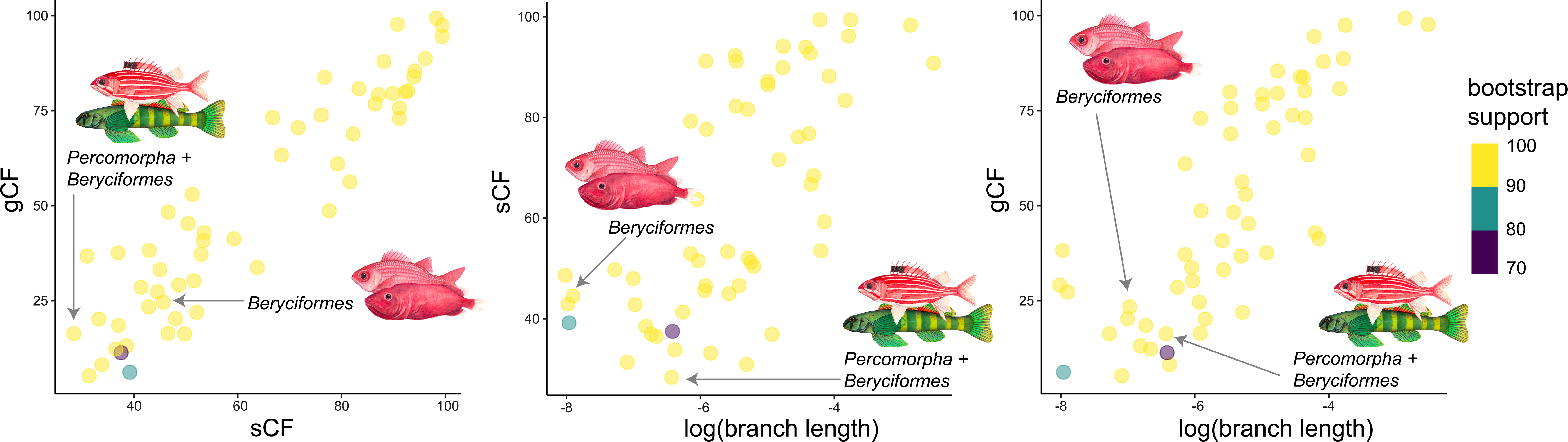
Phylogenetic Uncertainty Surrounding Acanthopterygian Relationships. Graphs show relationships between gene concordance factors (gCF), site concordance factors (sCF), and log-transformed branch lengths. Key nodes are indicated. Illustrations are by Julie Johnson.

Within *Trachichthyiformes*, *Diretmidae* (deep-sea spinyfins) are resolved as the sister lineage of all other species in the clade, the epipelagic *Anomalopidae* (flashlightfishes) and *Monocentridae* (porcupinefishes) form a clade, and *Anoplogaster* (fangtooths) and *Trachichthyidae* are sister lineages (Figure 1; Figures S1-S3, S5-S7. The phylogeny inferred from the Sanger-sequenced nuclear genes resolves *Trachichthyidae* as paraphyletic because *Anoplogaster* is placed as the sister taxon of a clade containing *Trachichthys* and *Aulotrachichthys*, albeit with weak nodal support (Figure S5). In the epipelagic *Holocentridae*, the *Myripristinae* (soldierfishes) and *Holocentrinae* (squirrelfishes) are each resolved as monophyletic lineages; however, the two sampled species of *Holocentrus* are nested in a paraphyletic *Neoniphon* (Figure 1; Figures S1-S3, S5-S7). Relationships among species of Holocentrinae vary among molecular studies and are characterized by relatively low node support (Dornburg et al. 2012; Vella et al. 2016; Rabosky et al. 2018; Deef 2021; Tang et al. 2023).

There are two inclusive clades of *Berycoidei*. *Melamphaidae* (ridgeheads) is nested in a paraphyletic *Berycidae* (alfonsinos) and both *Beryx* and *Centroberyx* are paraphyletic (Figure 1; Figures S1-S3, S5-S7). The other clade of *Berycoidei* consists of deep-sea species. *Barbourisia rufa* (Velvet Whalefish), *Stephanoberycidae* (pricklefishes) and *Cetomimidae* (flabby whalefishes) form a clade that is the sister lineage of a group containing *Rondeletia* (redmouth whalefishes), *Gibberichthys* (gibberfishes), and *Hispidoberyx ambagiosus* (Bristlyskin) (Figure 1; Figures S6-S7). Previous phylogenetic analyses using morphological characters resolves a monophyletic group containing *Hispidoberyx* and *Stephanoberycidae* or *Hispidoberyx* as the sister lineage of a clade that includes *Stephanoberycidae*, *Gibberichthys*, and *Rondeletia* (Kimura 2024). Ours is the first molecular study to include *Hispidoberyx. Gibberichthys* and *Rondeletia* are sister taxa in the only other molecular analyses that include the two lineages (Weber 2020; New Findings of the Rare Species Rondeletia bicolor (Stephanoberycoidei) Over the Mid-Atlantic Ridge and Some Peculiarities of the Rondeletiidae Family’s Phylologeny | Journal of Ichthyology n.d.). Phylogenetic relationships among sampled species of *Cetomimidae* are largely congruent with previous inferences based on morphology (Paxton 1989), mitochondrial DNA (Johnson et al. 2009; Kobyliansky et al. 2023), and UCEs (Ghezelayagh et al. 2022): *Procetichthys kreffti* is the sister lineage all other species in the clade, *Cetostoma regani* and *Ditropichthys storeri* are sister species, and species of *Cetomimus* are nested within a paraphyletic *Gyrinomimus* (Figure 1, Figures S1-S7).

### (b) The Timescale and Mode of Acanthopterygian Diversification

Our time-calibrated phylogeny estimates that time of most recent common ancestry for *Berycoidei, Holocentridae*, and *Trachichthyiformes* occurs in the Cenozoic, confirming that the species diversity of these clades results from divergence events restricted to the last 66 million years of Earth history (Figure 1). The most recent common ancestor (MRCA) of *Trachichthyiformes* and all other acanthopterygians dates to the Early Cretaceous, 105.5 Ma (95% highest posterior density, HPD, interval: 90.73, 124.81 Ma), but the crown clade originates in the Eocene at 48.5 Ma (95% HPD: 33.94, 72.81 Ma). Similarly, whereas *Beryciformes* and *Percomorpha* are estimated to last share common ancestry 98.2 million years ago (95% HPD: 86.75, 112.05 Ma), the age of the MRCA of crown lineage *Beryciformes* is estimated at 73.55 Ma (95% HPD: 66.4, 81.05 Ma), and the ages of the MRCAs of the living lineages of both *Holocentridae* (median MRCA age: 52.77 Ma, 95% HPD: 47.78, 57.97 Ma) and *Berycoidei* (median MRCA age: 60.22 Ma, 95% HPD: 43.33, 73.12 Ma) are dated to the Paleocene-Eocene. In *Trachichthyiformes*, all family-level crown clades originate in the Oligocene-Miocene, as do both subfamilies in *Holocentridae* and all family-level clades in *Berycoidei* except for *Cetomimidae* and *Melamphaidae*, which originate in the 7 million years following the Grande Coupere (Figure 1). In *Holocentridae*, species of *Myripristinae*, *Sargocentron*, and a clade consisting of *Flammeo, Neoniphon,* and *Holocentrus* show rapid successive divergences beginning at 20 million years ago.

The recent ages of many of the family-level crown clades across the clades most closely related to *Percomorpha* remove the relevance of the ecological preferences of living species in these lineages for understanding the deep-time evolution of acanthomorph fishes. This is observable in our ancestral state reconstruction of habitat preference, which reconstructs all deep nodes in our phylogeny as epipelagic except for the MRCA of *Berycoidei* (Figure 1); this broadly matches with expectations from the fossil record, where early acanthomorphs are abundant in shallow-marine and nearshore settings (Patterson and Patterson 1967; Delbarre et al. 2016; Andrews et al. 2023; Friedman et al. 2023). Secondly, the recent ages of deep-sea clades with potential specializations for life in the bathypelagic zone, such as extreme sexual dimorphism in *Cetomimidae* (Johnson et al. 2009) and pelagic cnidarian-mimicking larvae in cetomimids and *Gibberichthys* (Ho et al. 2023), indicates recent origins of these evolutionary novelties relative to the deepest divergences among the major lineages of *Acanthomorpha*. We find that reef association, sexual dimorphism, and bathypelagic habitat preference have differentially driven the diversification of clades like *Holocentridae* and *Cetomimidae* into shallow-water and deep-sea habitats, respectively (Figure 3A-C, E-F). In contrast, mesopelagic habitat preference and larval mimicry are not associated with increased rates of lineage diversification (Figure 3A-D).

**Figure 3.**
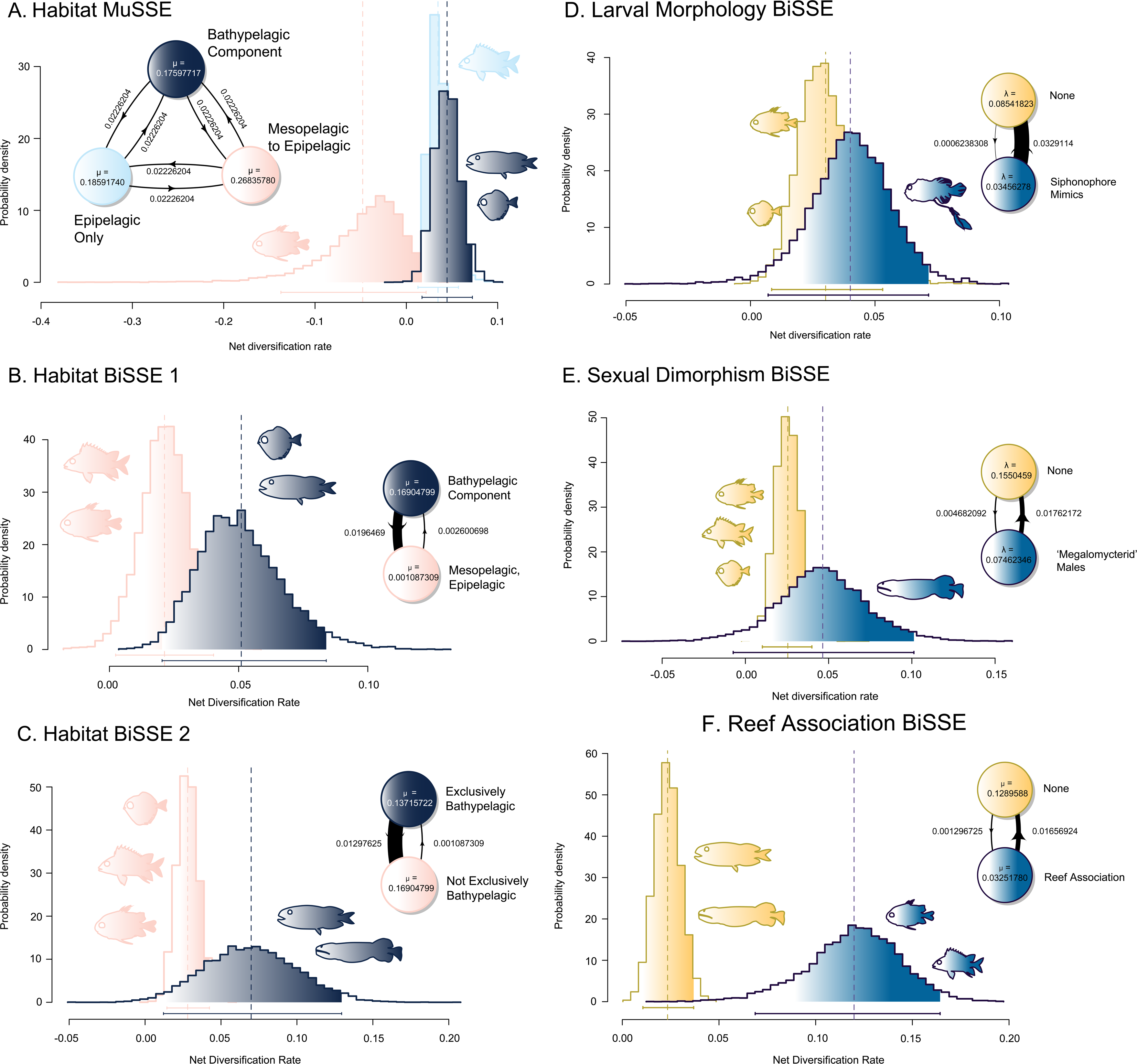
Factors Affecting the Diversification of Early-Diverging Acanthopterygians. Plots of posterior distributions of trait-associated diversification rates from BiSSE (A-B, D-F) and MuSSE (C) analyses of (A-C) habitat, (D) larval morphology, (E) sexual dimorphism, and (F) reef association. State transition diagrams indicate parameters allowed to vary in favored models (lambda = speciation, mu = extinction), and transition bars are bolded relative to their values. Silhouettes are by the author, made from public-domain images on phylopic.org and Wikimedia commons.

## Discussion

The phylogeny and timescale of evolution among the major lineages of *Acanthopterygii* presented in this study clarifies how the evolutionary history of *Berycoidei, Holocentridae*, and *Trachichthyiformes* illuminates our understanding of the origins of the exceptional diversity of spiny-rayed fishes. Despite their morphological similarities to some of the oldest fossils identifiable as crown acanthomorphs such as Cretaceous ‘berycid’ and ‘trachichthyiform’ otoliths and holomorphic body fossils (Patterson and Patterson 1967; Stewart 1996; Patterson and White 1997; Schwarzhans et al. 2017; Stringer et al. 2020), our divergence time estimates do not support Mesozoic origins for the living lineages of roughies, squirrelfishes, whalefishes, and relatives. Instead, all three of the crown clades that represent the closest relatives of *Percomorpha* have post-Cretaceous origins, and the shallow-water and deep-sea clades nested within them are recent diversifications that have taken place in the last 10 to 30 million years of Earth history (Figure 1). The ages for these clades inferred in this study suggest a reevaluation of the putative Cretaceous record of early beryciforms and trachichthyiforms is needed to resolve the affinities of these important early acanthomorph records (Andrews et al. 2023). The most charitable interpretation of these fossils using our time-calibrated phylogeny would place them on the stems of *Trachichthyiformes* and *Beryciformes* (sensu (Near and Thacker 2024)), implying convergent evolution in skeletal and otolith morphology between Mesozoic species and living forms nested deep within the crown clades.

Two events, the Grande Coupere (=Eocene-Oligocene Extinction)(Eocene-Oligocene Climatic and Biotic Evolution 1992; Prothero 1994; Kelley and Hansen 1996; Sibert et al. 2020) and the Early Miocene reef reorganization and turnover (Renema et al. 2008; DiBattista et al. 2018; Siqueira et al. 2023; Tian et al. 2024), are associated with the origination and rapid diversification of holocentrids, berycoids, and trachichthyiforms (Figure 1) into epipelagic reef and bathypelagic ecosystems (Figure 3). Among these, rapid diversification has occurred recently in *Melamphaes* and *Scopelogadus.* Other deep-sea clades, including *Anoplogaster*, *Cetomimidae*, and *Melamphaidae*, originated directly following the Grande Coupere (Figure 1). No clade that exclusively inhabits deep sea habitats appears to predate this extinction event, suggesting a recent assembly of deep-sea diversity within the most deeply divergent acanthopterygian lineages. This result implies recent origins for apparent adaptations to life in the mesopelagic and bathypelagic zones found across *Trachichthyiformes* and *Berycoidei*, such as the expanded opsin repertoire of spinyfins (Musilova et al. 2019) and the expanded sensory pore system and extraordinary sexual dimorphism of cetomimid whalefishes (Paxton 1989; Johnson et al. 2009), are recent innovations rather than relics of early acanthomorph radiations in the marine realm.

The ages of deep-sea invasions within early-diverging acanthopterygian clades match or slightly postdate inferred times for key deep-sea percomorph radiations (Betancur-R et al. 2013; Near et al. 2013; Alfaro et al. 2018; Hughes et al. 2018; Hotaling et al. 2021; Ghezelayagh et al. 2022), particularly ceratioid anglerfishes (Miller et al. 2023; Brownstein et al. 2024b), and highlights the Eocene-Oligocene as a key period of change in deep-sea vertebrate faunas characterized by the invasion of this nutrient-poor environment by several clades of acanthomorphs. This interval, which was characterized by extreme shifts in global climate (Eocene-Oligocene Climatic and Biotic Evolution 1992; Prothero 1994; Liu et al. 2009; Pearson et al. 2009; Hutchinson et al. 2021) that included rapid changes in oceanic temperature throughout the water column (Meckler et al. 2022), may have driven the differential dispersal and subsequent radiation of acanthomorph fishes, including berycoids and trachichthyiforms, into deep sea environments.

In contrast, the living diversity of holocentrids appears to have accumulated from successive diversification events that took place starting approximately 20 million years ago (Figure 1). Holocentrid diversification has previously been tied to the formation of the Indo-Pacific Biodiversity Hotspot . Our revised inference of the timescale of holocentrid evolution, which places most divergences between 2 and 5 million years later in time than previously estimated (Dornburg et al. 2015), remains congruent with this hypothesis (Renema et al. 2008; Cowman and Bellwood 2011, 2013; DiBattista et al. 2018; Tian et al. 2024).

The evolutionary history of *Percomorpha* and the most closely related clades of spiny-rayed fishes provides a strong example of how outgroups to a species-rich clade might not be ideal for inferring the ancestral habitats and phenotypes prefiguring such high diversity. Despite resembling fossils from the Mesozoic (Patterson and Patterson 1967; Patterson and White 1997; Friedman et al. 2023), living species in *Trachichthyiformes* and *Beryciformes* are the result of Cenozoic radiations into reef and bathypelagic habitats that have largely taken place in the last 30 million years (Figure 1). Our time-calibrated phylogeny indicates that iconic deep-sea fishes, such as the cetomimid whalefishes and fangtooths, invaded the deep sea during the Eocene to Oligocene Epochs of the Paleogene, a period characterized by rapid global sea temperature fluctuations (Figure 1; Figure 3A-C). Further, we tie the evolution of reef-associated clades like holocentrids, anomalopids, and monocentrids to reef reorganization and hotspot movement during the Miocene (Figure 1, Figure 3F). Together, these results show that the evolutionary history of *Beryciformes* and *Trachichthyiformes* has been shaped by global climate events and biotic turnovers that far postdate their most recent common ancestry with other lineages of acanthopterygian fishes. Rather than representing holdovers from an ancient acanthomorph diversification, the present diversity of fishes in *Berycioidei, Holocentridae*, and *Trachichthyiformes* has been shaped by Cenozoic Earth system processes that affected most other marine acanthomorph clades (Renema et al. 2008; Cowman and Bellwood 2011, 2013; Dornburg et al. 2015; Arcila and Tyler 2017; DiBattista et al. 2018; Sibert et al. 2020; Miller et al. 2022; Brownstein et al. 2024a).

## Data Availability

All data relevant to the results presented in this manuscript is included in the online supplement to this article or deposited on Dryad. Newly-generated nuclear gene sequences are on the Dryad repository and will be deposited on the NCBI repository GenBank. **Acknowledgements.** We thank members of the Near and Dornburg labs for discussions regarding this manuscript. CDB thanks Spencer Lott for guidance on processing genomes using phyluce. We also thank G.D. Johnson, J.A. Morre, and the late J.R. Paxton for tissues. Additional tissues were provided by the Biodiversity Research Museum, Academia Sinica, Taiwan.

## Fossil Calibration Justification List

### †Miobarbourisia aomori

**Phylogenetic Placement and Justification:** †*Miobarbourisia aomori* calibrates the split between *Barbourisia rufa* and *Acanthochaenus luetkenii*, and is identified as a pan-*Barbourisia* based on the presence of the following combination of features (from Fujii et al. [1]): large subterminal mouth and jaw that extends posterior to small eyes, an expanded sensory canal system on the head; enlarged scales with small pores run parallel to the enlarged canal on the body; diminutive scalation across the body; no dorsal, pelvic, or anal fin spines; pelvic fins on the abdomen and with six rays; prominent, well-separated procurrent caudal rays; triangular, ridged opercle; elongated, flattened gill rakers.

**Stratigraphy:** Neogene Taga and Hitachi groups in the Kitaibaraki-Takahagi area, Ibaraki Prefecture, Japan: Middle to Late Miocene sedimentary complexes of shelf to slope deposits, submarine channel fills and submarine slide scar fills [1].

**Fossil age**: 7.5 Ma.

### †Eoholocentrum macrocephalum

**Phylogenetic Placement and Justification:** †*Eoholocentrum macrocephalum* calibrates the MRCA of crown *Holocentridae* and is placed there following the results of a recent phylogenetic analysis of morphological characters [2].

**Stratigraphy:** Monte Bolca Lagerstätten, Italy: Ypresian, Eocene [3].

**Fossil age**: 48.5 Ma.

### †Africentrum melitense

**Phylogenetic Placement and Justification:** †*Africentrum melitense* calibrates the MRCA of crown *Myripristinae* and is placed there following the results of a recent phylogenetic analysis of morphological characters [2].

**Stratigraphy:** Pietra de Cantoni, Malta: Ypresian, Early Miocene, 17.154 Ma [2].

**Fossil age**: 17.154 Ma

### †Iridopristis parrisi

**Phylogenetic Placement and Justification:** †*Iridopristis parrisi* calibrates the MRCA of *Berycioidei* and *Holocentridae* and is placed as a pan-holocentrid in a recent phylogenetic analysis of morphological characters [2].

**Stratigraphy:** Main Fossiliferous Layer, Hornerstown Formation, New Jersey, USA: earliest Danian, Paleocene [2], approximately 66.02 Ma [4].

**Fossil age**: 66.02 Ma.

### †Scopelogadus sp

**Phylogenetic Placement and Justification:** †*Scopelogadus* sp. calibrates the MRCA of *Scopelogadus* and *Melamphaes*, and is placed as a member of pan-*Scopelogadus* among melamphaeids based on fin ray and vertebral counts and the presence of cycloid scales [5]. **Stratigraphy:** Kurasi Formation of Sakhalin Island, Russia: Late Miocene [5].

**Fossil age**: 5.333 Ma.

### †Gephyroberyx robustus

**Phylogenetic Placement and Justification:** †*Gephyroberyx robustus* calibrates the MRCA of *Gephyroberyx* and *Hoplostethus*. See Near et al. [6] and Ghezelayagh et al. [7] for a full justification.

**Stratigraphy:** Oligocene of the Caucasus, Russia: Rupelian, 33.9 to 27.82 Ma [8].

**Fossil age**: 27.82 Ma.

### †Cumbaaichthys oxyrhynchus

**Phylogenetic Placement and Justification:** †*Cumbaaichthys oxyrhynchus* calibrates pan- *Polymixia*; in analyses presented in this paper, this is the MRCA of *Polymixia lowei* and *Percopsiformes*. See a full justification of this calibration in [9].

**Stratigraphy**: Lacdes Bois, Northwest Territories, Canada: early Turonian, Late Cretaceous [9].

**Fossil tip age**: 93.9 Ma (M. Wilson 1977; Mayr et al. 2019). CHECK AGES

### †Trichophanes foliarum

**Phylogenetic Placement and Justification:** †*Trichophanes foliarum* calibrates pan-

*Aphredoderus*; in analyses presented in this paper, this is the MRCA of *Aphredoderus sayanus* and *Percopsis omiscomaycus*. This placement is based on a phylogenetic analysis of morphological characters [10]. See a full justification of this calibration in [7,8].

**Stratigraphy**: Florissant Fossil Beds, Colorado, USA: Priabonian, Eocene 37.7-33.9 Ma [8].

**Fossil tip age**: 35.8 Ma.

### †Gasterorhamphosus zuppichini

**Phylogenetic Placement and Justification:** †*Gasterorhamphosus zuppichini* calibrates the MRCA of *Thunnus albacares* and all percomorphs except *Carapus bermudensis* and *Batrachoides pacifici*. †*G. zuppichini* is the oldest identifiable member of *Syngnathiformes* and is placed in the crown of that order based on the results of phylogenetic analyses of morphological characters [11].

**Stratigraphy**: Nardo, Italy; Late Cretaceous, Santonian-Campanian, 83.6 Ma [11].

**Fossil tip age**: 83.6 Ma.

### †Ctenoplectus williamsi

**Phylogenetic Placement and Justification:** †*Ctenoplectus williamsi* is a member of *Tetraodontoidei* in *Acanthuriformes* sensu [7,8] and is placed in that clade based on the results of phylogenetic analysis of morphological characters [12]. In the analysis present in this paper, †*Ctenoplectus williamsi* calibrates the split between the tetraodontoid-lophioid clade (represented by *Sladenia gardineri*) and *Dicentrarchus labrax*.

**Stratigraphy**: London Clay Formation, England, UK: Ypresian, Eocene, Paleogene [12].

**Fossil tip age**: 53.0 Ma.

### †Mene purydi

**Phylogenetic Placement and Justification:** †*Mene purydi* is assignable to pan-*Mene* and calibrates the MRCA of *Betta prima* and *Caranx caninus* in our phylogeny. See a full justification of this calibration in [7,8].

**Stratigraphy**: Talara Province, northwestern Perú. Thanetian to Ypresian, Eocene [13], minimum 56.0 Ma [4].

**Fossil tip age**: 56.0 Ma.

### †Paleoserranus lakamhae

**Phylogenetic Placement and Justification:** †*Paleoserranus lakamhae* is a probable crown percomorph [14](“*Serranidae*” incertae sedis; this family is paraphyletic [7]) and calibrates the MRCA of *Percina burtoni* and *Zoarces andriashevi* in analyses presented in this paper. This species is allied with “*Serranidae*” based on the absence of a posterior uroneural and procurrent spur on the ventral caudal fin [14].

**Stratigraphy**: División del Norte and Belisario Domínguez quarries, Tenejapa-Lacandón Formations, Chiapas, Mexico; Danian, Paleocene [14].

**Fossil tip age**: 61.6 Ma [14].

## Supplementary Figure Captions

**Figure S1.**
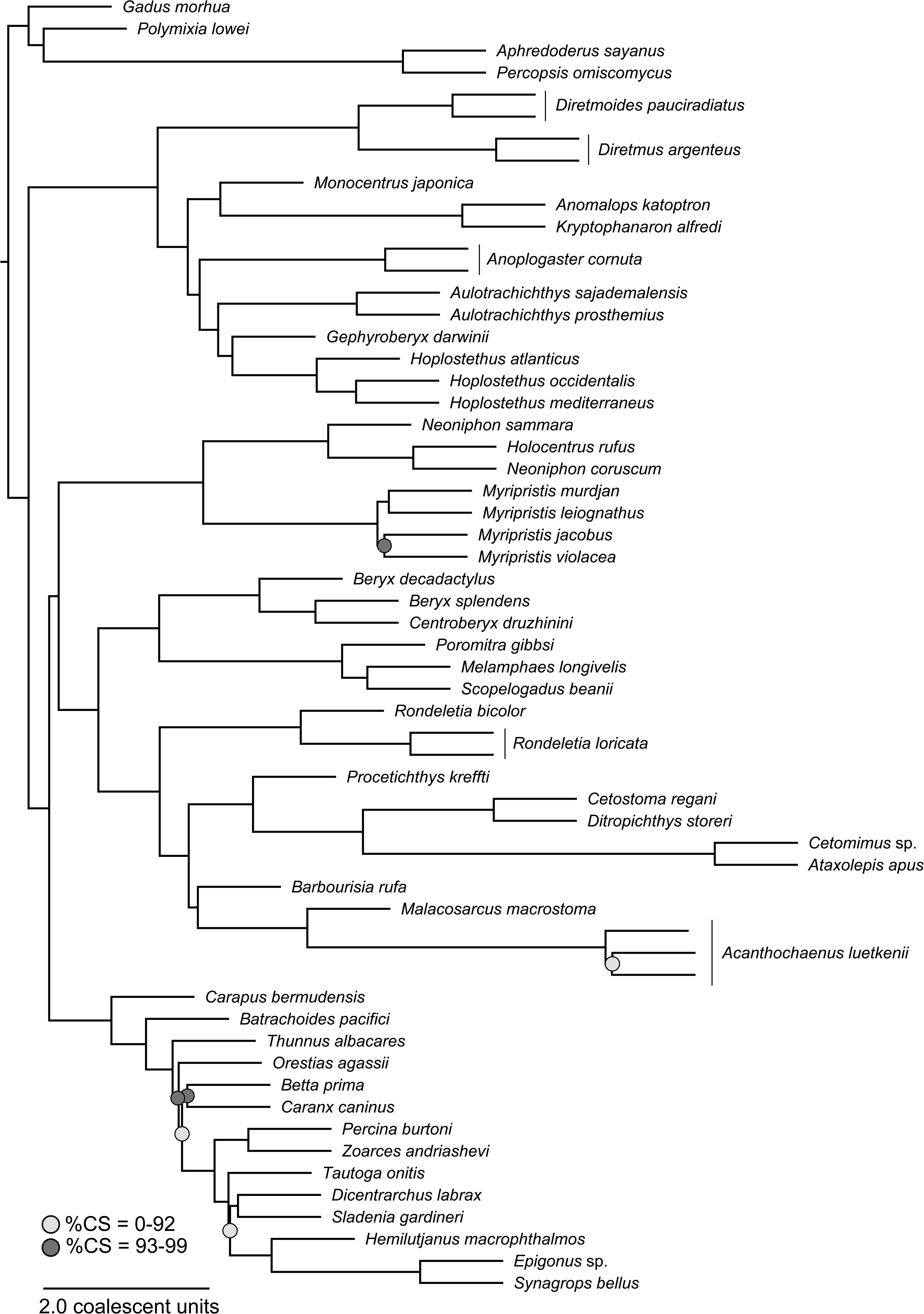
ASTRAL-III phylogeny inferred using only the 879 UCE sequences. %CS = percent coalescent support.

**Figure S2.**
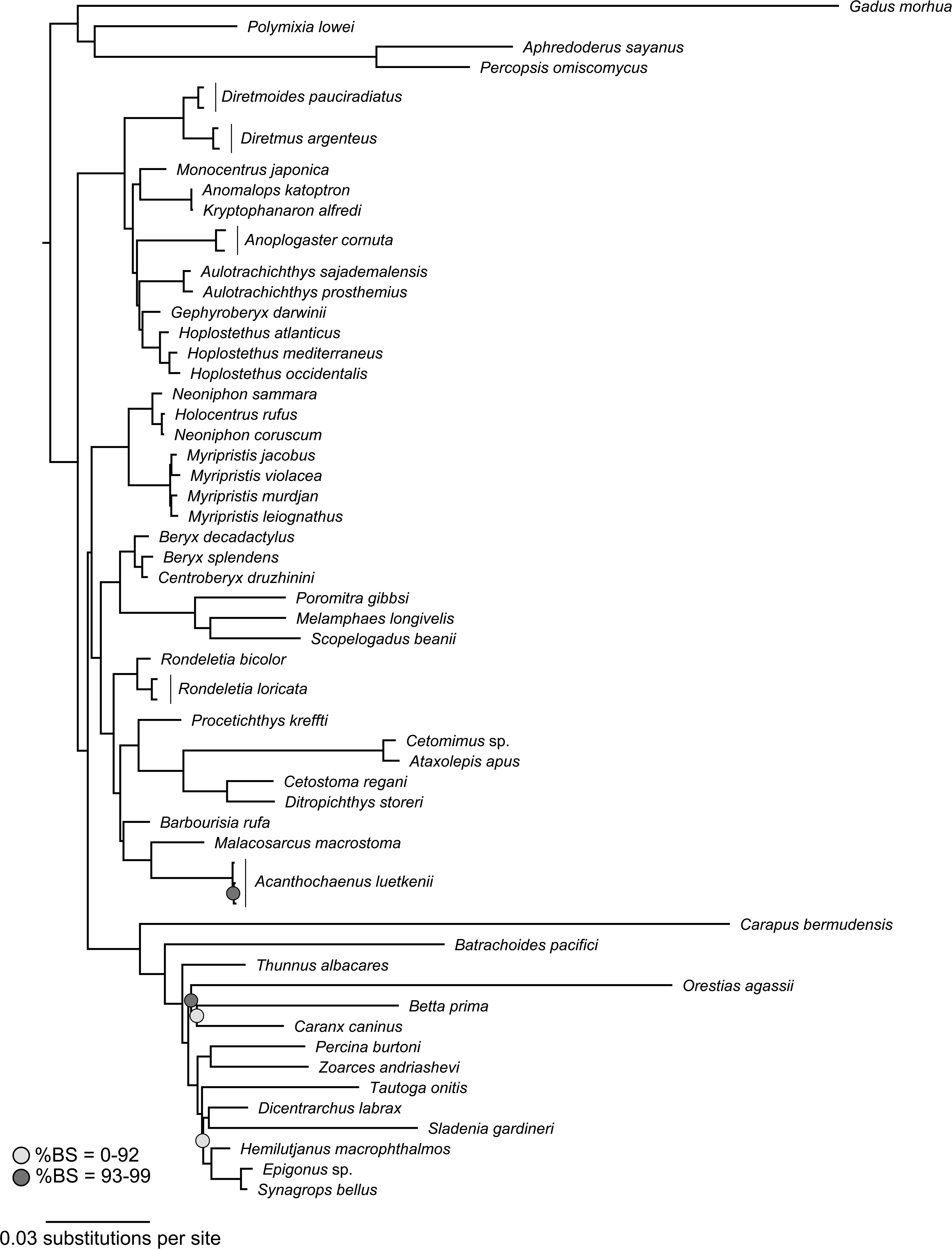
Phylogeny inferred using only the concatenated 879 UCE sequences analyzed as a single partition. %BS = percent ultrafast bootstrap support.

**Figure S3.**
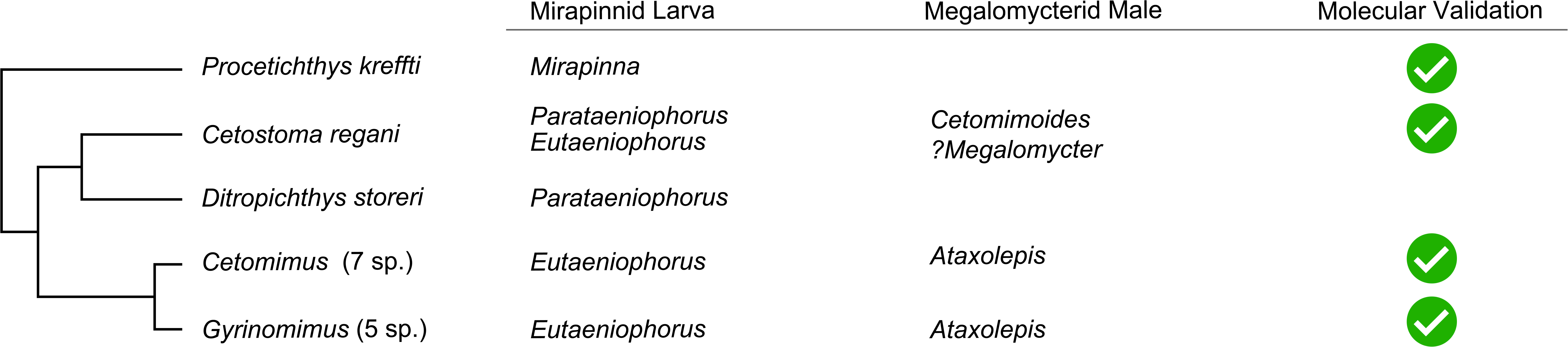
Phylogeny inferred using only the concatenated 879 UCE sequences analyzed as multiple partitions. %BS = percent ultrafast bootstrap support.

**Figure S4.**
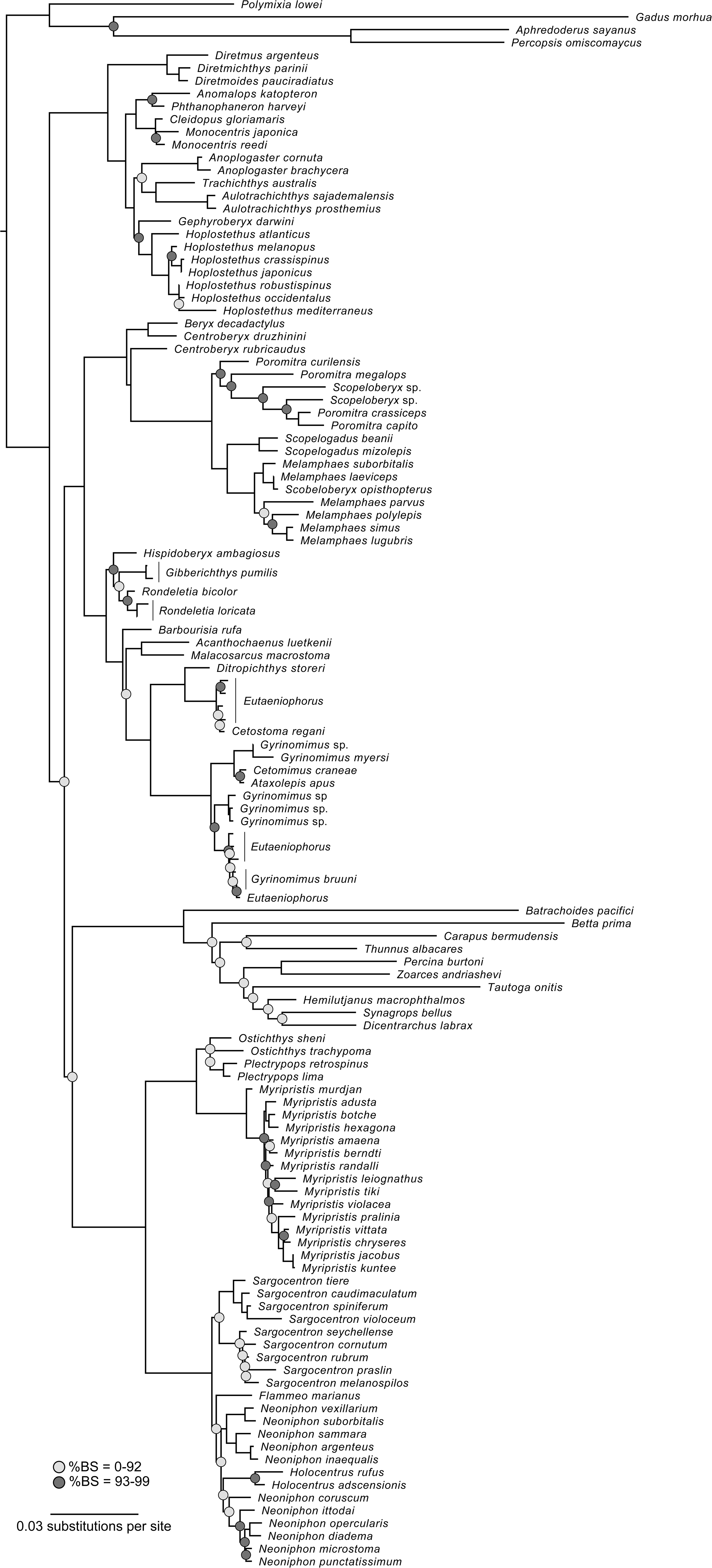
Correspondence of ‘megalomycterid’ males and ‘mirapinnid’ larvae to cetomimid species, based on the results of phylogenetic analysis of Sanger sequences employed in this study, barcoding from Ghezelayagh et al. [7], and mitochondrial DNA-based phylogenies [15,16].

**Figure S5.**
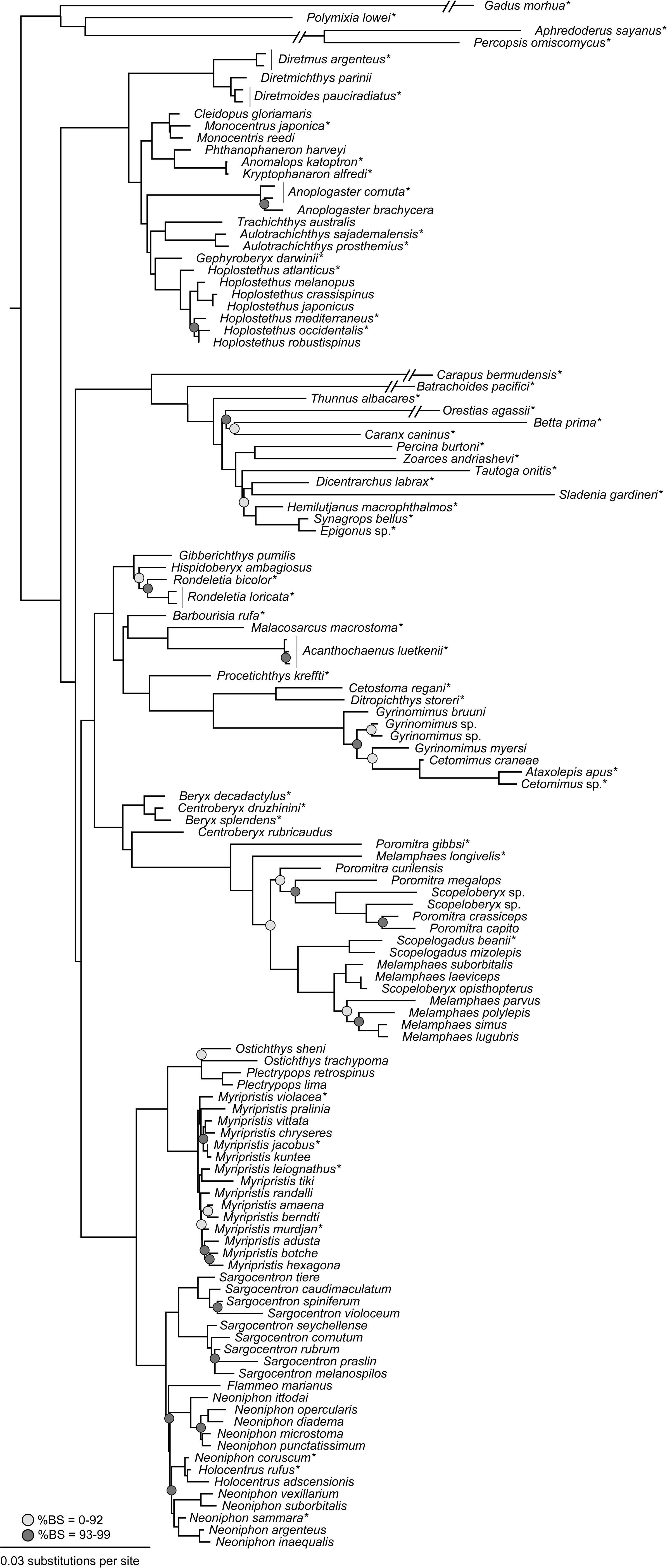
Phylogeny inferred using only the concatenated Sanger-sequenced nuclear gene sequences analyzed as a single partition. %BS = percent ultrafast bootstrap support.

**Figure S6.**
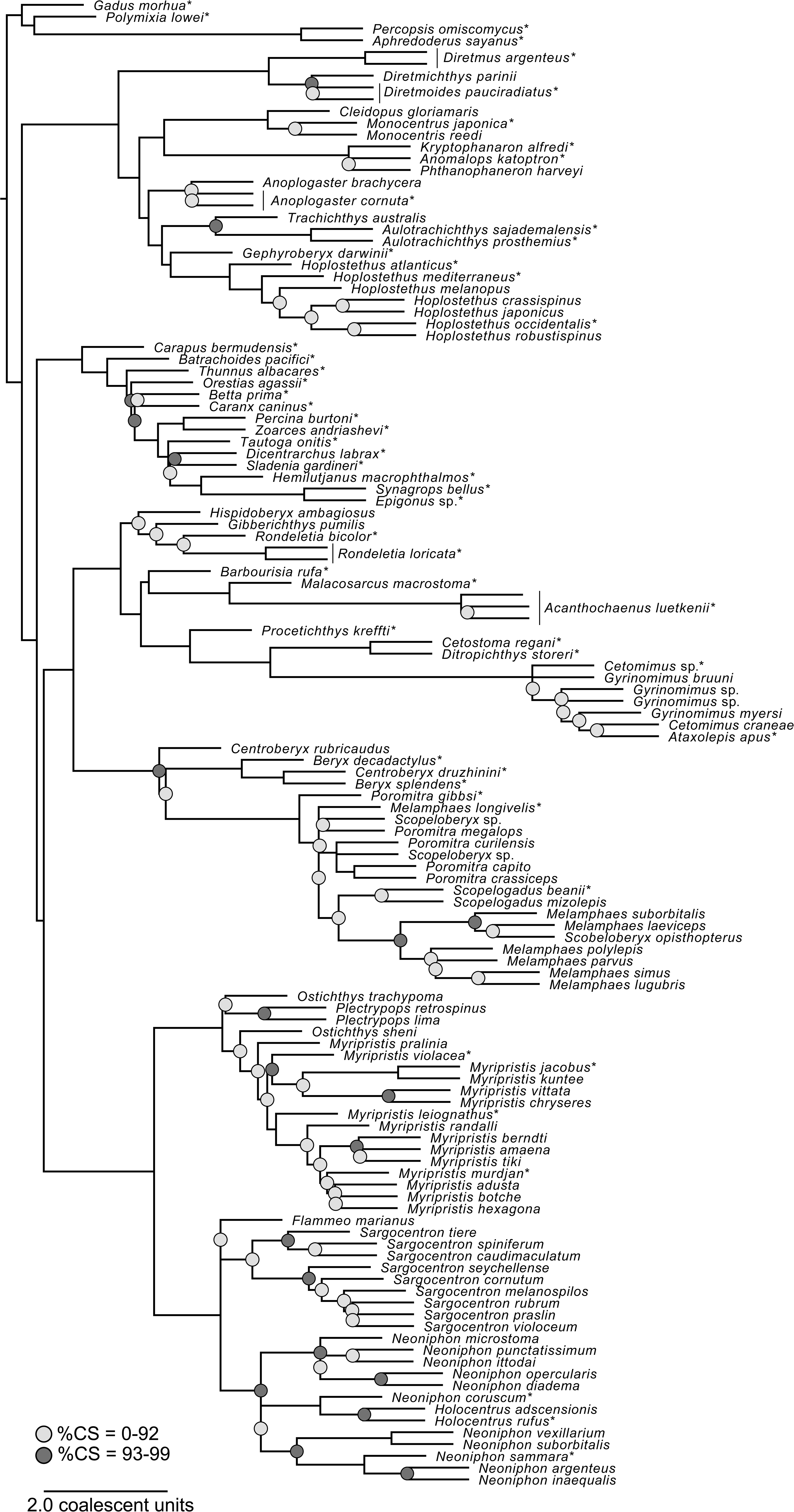
Phylogeny inferred using the concatenated 879 UCE sequences and Sanger-sequenced nuclear gene sequences analyzed as a single partition. %BS = percent ultrafast bootstrap support.

**Figure S7.**
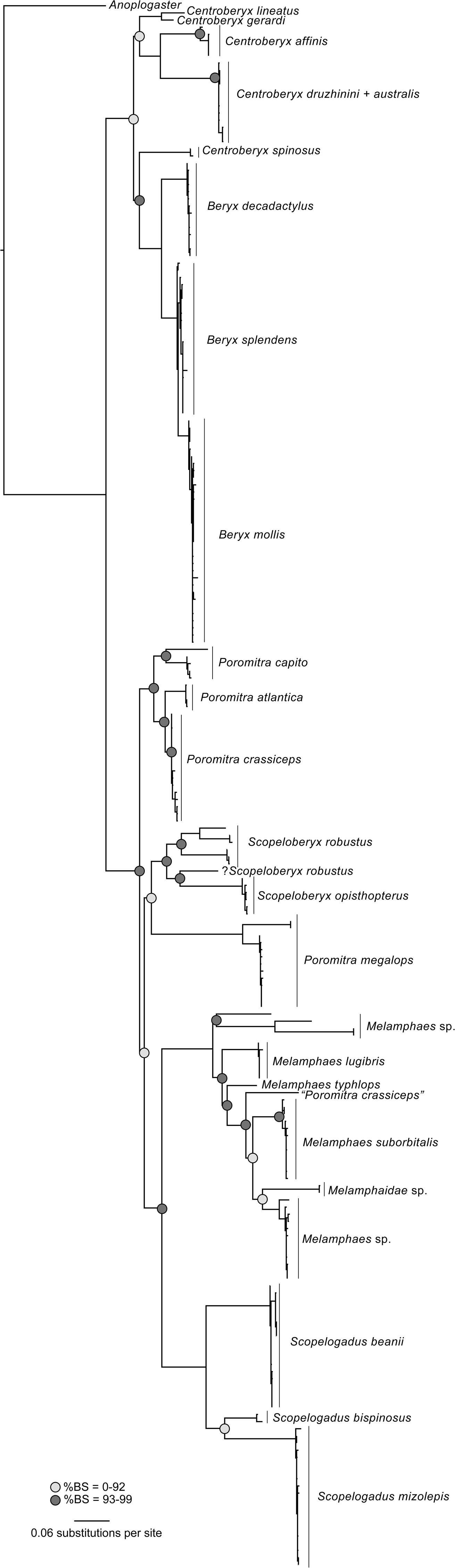
ASTRAL-III phylogeny inferred using the 879 UCE sequences and Sanger-sequenced nuclear gene sequences. %CS = percent coalescent support.

